# Microbial priming controls terrestrial carbon burial in marine sediments

**DOI:** 10.64898/2026.10.21.760051

**Authors:** Josefine Friederike Weiß, Kathleen R. Stoof-Leichsenring, Christiane Boeckel, Lester Lembke-Jene, Vera D. Meyer, Uwe John, Ulrike Herzschuh

## Abstract

Marine sediments are a globally important sink for organic carbon of both marine and terrestrial origin, yet far less terrestrial carbon is ultimately buried than rivers deliver to the ocean margins. This long-standing “terrestrial carbon conundrum” has resisted explanation, largely because conventional bulk proxies cannot distinguish marine, terrestrial and microbial-degrader carbon pools. Here we use sedimentary ancient DNA from five records spanning the Bering Sea to the Sea of Okhotsk over the past 30,000 years to reconstruct organic carbon resolved simultaneously by taxonomy and habitat. Integrating these taxon-resolved carbon estimates with environmental proxies across the last glacial–interglacial cycle, we find that terrestrial carbon burial is governed not only by supply but by biogeochemical interactions previously documented only in soils. Microbial processing splits into two competing pathways: under priming, marine organic matter fuels the degradation of terrestrial organic matter, whereas under preferential substrate utilization microbes consume marine organic matter and spare terrestrial carbon. Both pathways are triggered only once marine organic-matter availability crosses a critical, temperature-governed threshold. This temperature-sensitive cascade reveals a previously unrecognized vulnerability of the ocean’s carbon sink to global warming.

## Introduction

Marine sediments represent the largest long-term reservoirs of organic carbon on Earth, playing a critical role in regulating the global carbon cycle and atmospheric CO_2_ levels over geological timescales (Burdige, 2007). A considerable fraction of this buried sedimentary organic matter is of terrestrial origin, delivered mainly by rivers and coastal erosion (Regnier et al., 2022). Terrestrial organic matter has traditionally been regarded as refractory (i.e. highly resistant to degradation) and therefore prone to efficient burial and storage in marine sediments, owing to macromolecular constituents such as lignin (Hedges et al., 1997). However, only one third of the transported terrestrial organic matter is actually found in marine sediments, a mismatch referred to as the terrestrial carbon conundrum (Bianchi, 2011).

Numerous studies have addressed the discrepancy between the amount of terrestrial organic matter delivered to the ocean by rivers and the amount ultimately buried in marine sediments (Bianchi, 2011; Goranov et al., 2025; Yi et al., 2025). Proposed explanations include oxidative camouflage masking the terrestrial signature through chemical and isotopic transformations that make the remaining terrestrial organic matter appear entirely marine-like (Goranov et al., 2025). Further studies point to analytical limitations in source identification (Yi et al., 2025), and rapid remineralization in the coastal ocean supported by microbes (Bianchi, 2011; Blair & Aller, 2012; Burd et al., 2016; Kandasamy & Nagender Nath, 2016; Jiao et al., 2024) as the main reasons for the discrepancy. The mechanisms driving these high remineralization rates remain discussed, given the refractory character of terrestrial organic matter (Blair & Aller, 2012). Organic matter degradation by microbes is a complex process controlled by factors such as redox conditions, mineral associations, and community-governed metabolic capacity (Chen et al., 2023). While some of these controls are comparatively well understood in terrestrial and freshwater ecosystems, they have received little attention in marine settings. Studies in soils and freshwater systems demonstrate, for example, that labile carbon inputs can trigger the degradation of refractory organic matter, an effect known as priming (Kuzyakov, 2010; Bianchi et al., 2015). Specific microbial communities can use the energy and nutrients derived from easy accessible (labile) substrates to synthesize extracellular enzymes that target more refractory compounds. As a result, fresh organic inputs can drive the remineralization of otherwise stable, refractory organic matter (Bianchi, 2011). Recent studies suggest that similar processes occur in marine environments (Sanches et al., 2021; Zhu et al., 2024), however, no consistent findings indicating an interaction between the marine and terrestrial carbon pool have been identified.

Detecting mechanisms like priming in the sedimentary record requires tracking both the terrestrial and marine organic matter sources and the microbial communities that consume them. Established geochemical proxies, such as stable carbon isotopes (□^13^C) and C/N ratios, provide reliable bulk estimates of terrestrial versus marine contributions (Riethdorf et al., 2013; Cao et al., 2023). Molecular markers like lignin phenols and long-chain *n*-alkanes (leaf waxes) offer even greater source specificity (Feng et al., 2013; Zhang et al., 2025). However, these latter purely geochemical approaches only analyze fractions of the carbon pool. The limitations of conventional proxies in simultaneously resolving terrestrial inputs, marine production, and the specific degrader community leave a critical gap in reconstructing long-term carbon dynamics.

Sedimentary ancient DNA (sedaDNA) together with shotgun metagenomics offer a promising solution to this gap, providing high-resolution taxonomic profiles that encompass the entire tree of life found in marine sediments (Zimmermann et al., 2023, Weiß et al., 2025, Schreiber et al., 2025, Herzschuh et al., 2025, Boeckel et al., 2026). This approach facilitates the identification of both source organisms of organic matter and their potential microbial degraders, however, the lack of a method to translate sedaDNA into quantitative carbon estimates has prevented the analytical decoupling of terrestrial and marine sources yet. While the conversion of sedimentary ancient DNA into taxon-specific carbon estimates (genC) is beginning to close this gap (Yong & Weiß et al., 2026 preprint), its application has so far been limited to lacustrine settings.

Here, we bridge this quantitative gap to reconstruct the dynamics of terrestrial and marine carbon pools from the late Glacial through the Holocene. In this study, we apply shotgun metagenomics to five high-resolution sediment cores along a North Pacific transect to reconstruct reactive terrestrial and marine carbon pool dynamics. Through the application of the novel genC pipeline, we present the first semi-quantitative evidence that microbial community assembly and its interaction with marine primary production serve as fundamental drivers of terrestrial carbon degradation in the ocean. Ultimately, these findings demonstrate that integrating microbial ecology into geological carbon budgets and understanding paleoceanographic dynamics is essential to fully resolve the terrestrial carbon conundrum.

## Results and discussion

We generated a comprehensive metagenomic shotgun dataset comprising over 105 million reads (8.5 to 30.7 million reads per site) from five marine sediment cores, which form a north-south transect extending from the Bering Sea to the Okhotsk Sea (**Fig. 1A**; Table S1). These records span the transition from the late Glacial (~25 ka) through the Holocene, capturing major climatic shifts and associated dynamics in marine productivity and terrestrial influx (**Fig. 1B**).

**Figure 1.**
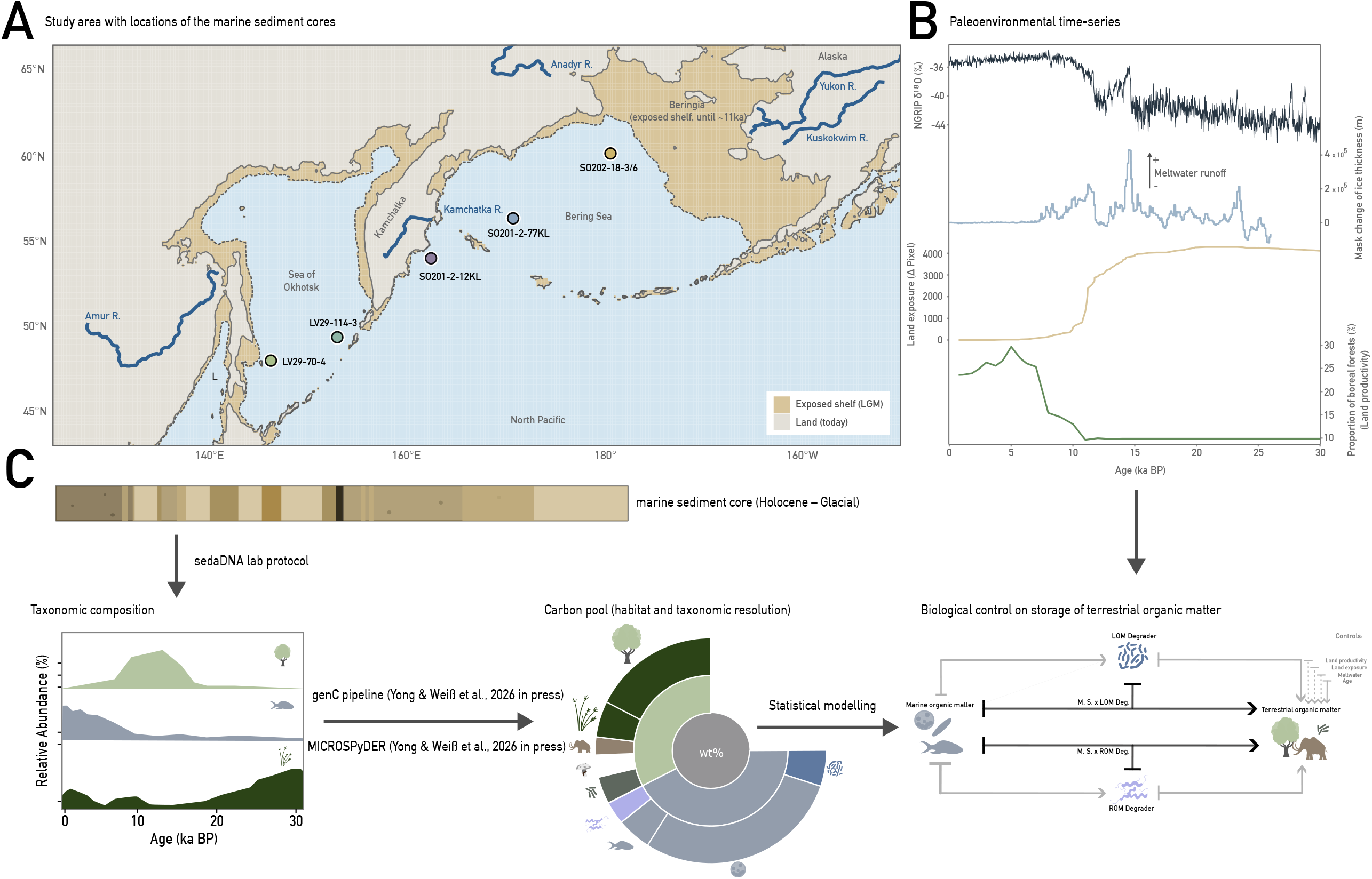
Study area, environmental context and analytical workflow. **A** Locations of the five subarctic North Pacific sediment cores (SO201-2-12KL, SO201-2-77KL, SO202-18-3/6, LV29-114-3, LV29-70-4) across the Bering Sea and Sea of Okhotsk, together with the major rivers delivering terrestrial organic matter (Amur, Anadyr, Yukon, Kuskokwim and Kamchatka). Shading distinguishes present-day land from the continental shelf exposed during the Last Glacial Maximum (Beringia) until ~11 ka BP. **B** Terrestrial-organic-matter delivery proxies over the last 30 kyr (meltwater flux, GLAC-1D (Peltier et al., 2015; Kapsch et al., 2022); land exposure (Herzschuh et al., 2025 preprint); boreal-forest proportion as a land-productivity proxy (Herzschuh et al., 2025 preprint)) and the NGRIP □^1^□O temperature record (Rasmussen et al., 2006), used as covariates in the model in C. C Analytical workflow. Sedimentary ancient DNA (sedaDNA) extracted from each core yields the down-core taxonomic composition (relative abundance), which is converted into semi-quantitative organic-carbon pools through the genC pipeline and MICROSPyDER habitat assignment (Yong & Weiß et al., 2026 in press). The resulting pools (shown at habitat and taxonomic resolution; central value, total organic carbon in wt %) are then analysed with a structural equation model of the biological control on terrestrial-organic-matter storage, in which marine organic matter interacts with labile-(LOM) and refractory-organic-matter (ROM) degraders, while controlling for land productivity, land exposure, meltwater and age.

Applying the genC pipeline to marine sediment cores for the first time (**Fig. 1C**), we found that the taxonomically resolved fraction contributes an average of 0.07–0.25 wt% to total organic carbon over all time intervals. Approximately half of the OC_DNA-projected_ (wt%) contribution (48–61% of the total OC_DNA-projected_ (wt%)) was assigned to a specific source habitat (marine or terrestrial) using MICROSPyDER (**Fig. 2A**). Marine taxa constituted the majority of this habitat-specific fraction, accounting for 31.8–44.8% across all five cores. Within this marine pool, primary producers were the dominant contributors to the organic matter (31.1–44.0%), while microbial degraders targeting rather refractory organic matter (ROM-degrader) and labile organic matter (LOM-degrader) accounted for only a minor fraction (0.04–2.4%). By comparison, terrestrial taxa accounted for a minor baseline contribution ranging from 3.9 to 14.0% (**Fig. 2A**). This aligns with previous bulk geochemical studies (Burdige, 2005; Tao et al., 2023). Integrating spatial and temporal dynamics, the reconstructed OC_DNA-projected_ (wt%) from all five cores across the North Pacific, tracks the relative organic matter of terrestrial (e.g. plants and terrestrial metazoa) and marine origin (e.g. phytoplankton and marine metazoa), alongside their potential microbial degraders (ROM– and LOM–degrader) over the past ~30 kyr (**Fig. 2B**).

**Figure 2.**
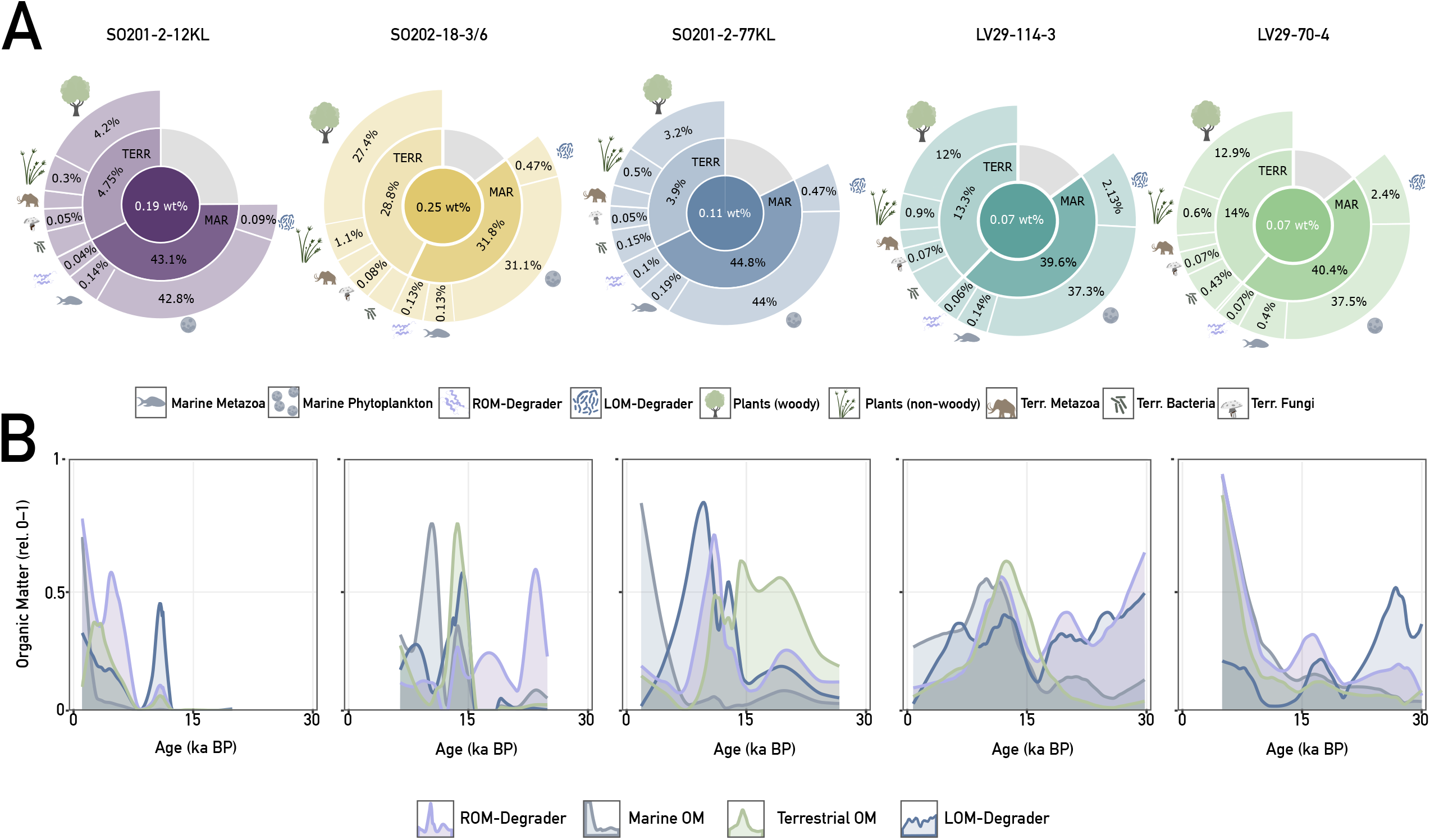
Sedimentary ancient DNA-based reconstruction of organic-carbon sources (genC) in the five subarctic North Pacific, Bering Sea and Sea of Okhotsk cores and over time. **A** Nested composition of DNA-projected organic carbon (genC) for each core (SO201-2-12KL, SO202-18-3/6, SO201-2-77KL, LV29-114-3, LV29-70-4). The central value is the mean total organic carbon reconstructed from DNA (wt % of dry sediment); the inner ring partitions it into marine (MAR) and terrestrial (TERR) sources, and the outer ring into taxon groups — marine metazoa, marine phytoplankton, refractory-organic-matter (ROM) degraders and labile-organic-matter (LOM) degraders (marine); woody plants, non-woody plants, terrestrial metazoa, bacteria and fungi (terrestrial). Values are the percentage contribution of each group to total genC (averaged over the core). Segment sizes are not proportional to these values; small segments were enlarged to ensure readability. **B** Down-core dynamics of the four key functional pools — marine organic matter, terrestrial biomass, ROM degraders and LOM degraders — shown as relative organic matter (each scaled 0–1) versus age (ka BP) for the same five cores. Curves are loess-smoothed.

The DNA-projected organic carbon is substantially smaller than bulk TOC, as expected, because a large and variable share of marine sedimentary TOC consists of black (pyrogenic) and petrogenic (fossil, rock-derived) carbon that is biologically inert and leaves no DNA signature (Meyer et al., 2019). We therefore validated genC not against bulk TOC but against source-specific proxies that each represent a defined organic-matter group, using n-alkanes for terrestrial input, opal for marine primary production, and brGDGTs for soil bacteria (Meyer et al. 2019; Max et al., 2012; Riethdorf et al., 2013). A Spearman rank analysis confirmed a significant positive correlation between the group specific genC-projected values and the established proxy records like biogenic opal, brGDGT, lignin and n-alkanes (**Supplementary Figure 1**).

Across all cores, marine organic matter was higher during the Holocene compared to glacial periods, a pattern widely supported by regional records (Riethdorf et al., 2013). Notably, while the deglacial showed only a minimal increase in cores SO201-2-12KL and SO202-18-3/6, core LV29-114-3 stands out as the only site where marine organic matter was higher during the deglacial than the Holocene (**Fig. 2B**). In contrast, terrestrial organic matter exhibits a more variable pattern over time. Whereas the more coastal sites (SO201-2-12KL, SO202-18-3/6, LV29-114-3, LV29-70-4) are characterized by pronounced peaks during the deglacial and Holocene, core SO201-2-77KL, situated further offshore, displays elevated terrestrial organic matter earlier in the record, spanning the Last Glacial Maximum into the early deglaciation (**Fig. 2B**), reflecting the closer proximity of terrestrial sources to this site during times of lower glacial sea levels (Lambeck et al., 2014).

Across three cores (SO201-2-12KL; SO202-18-3/6; SO201-2-77KL), the LOM- and ROM-degraders largely co-vary with the amounts of terrestrial and marine organic matter (**Fig. 2B**). While the amount of microbial degrader in the Sea of Okhotsk (LV29-114-3 & LV29-70-4) still tracks periods of high organic matter, this co-variation is less pronounced than in the more northern records. The concurrent increase of all organic matter and degrader pools highlights that these records overcome the closed-sum limitations inherent to traditional relative sedaDNA data.

### Marine organic matter fuels microbial degradation of terrestrial organic matter

The amount of terrestrial organic matter buried in marine sediments depends on two primary factors: how much enters the system, and the marine conditions it encounters, which determine how much of the organic matter is degraded prior to burial. To explore how the interplay of environmental and biological conditions shapes the carbon record in our North Pacific sediments, we developed a quantitative model of terrestrial organic matter dynamics and its drivers.

To systematically evaluate how strongly individual predictors shape terrestrial organic matter degradation or deposition across the sedimentary record, we quantified the coupling between organic matter and its degraders using a generalized linear mixed model (Gamma family, log link; marginal R^2^ = 0.73, conditional R^2^ = 0.81, **Supplementary Figure 2**). The response variable was terrestrial organic matter, and the predictors were marine organic matter as well as the biomass of the LOM- and ROM-degrader groups. Crucially, we included interaction terms between each degrader group and marine organic matter to evaluate whether the availability of marine carbon actively modifies the capacity of microbes to degrade terrestrial material. We included sample age and terrestrial-delivery covariates (land productivity, land exposure, meltwater) as controlling factors, and core identity as a random effect, to isolate distinct effects of microbial degradation.

The mixed model estimates the terms simultaneously but treats them as parallel predictors. This structure does not reflect the sequential biological pathway, where marine organic matter supports the degrader groups, which then act on the terrestrial organic matter pool. To make these dependencies explicit, we translated the fitted coefficients of the mixed model into a piecewise structural equation model (**Fig. 3A**). The piecewise structural equation model shows that marine organic matter increases the biomass of both degrader groups (substrate to LOM-degrader □ = +0.21, p < 0.01, substrate to ROM-degrader □ = +0.28, p < 0.01) (**Fig. 3A**). Furthermore, each degrader group affects terrestrial organic matter directly (LOM-degrader □ = +0.67, p < 0.001, ROM-degrader □ = +0.29, p < 0.05) and through its interaction with the marine substrate (**Fig. 3A**). These two interaction paths diverge in sign, positive for LOM degraders (standardized path □ = +1.98, p < 0.001, **Fig. 3A**, Gamma GLMM □ = +3.1, p < 10□□) and negative for ROM degraders (standardized path □ = −1.35, p < 0.001, Fig. 3A, Gamma GLMM □ = −2.0, p < 10□^3^). None of the terrestrial delivery proxies were significant within the model (land productivity □ = −0.20, land exposure □ = −0.05, meltwater □ = −0.01, all non-significant). Sample age, however, showed a significant negative effect, indicating that older samples contained less terrestrial organic matter (□ = −0.31, p < 0.001) (**Fig. 3A**).

**Figure 3.**
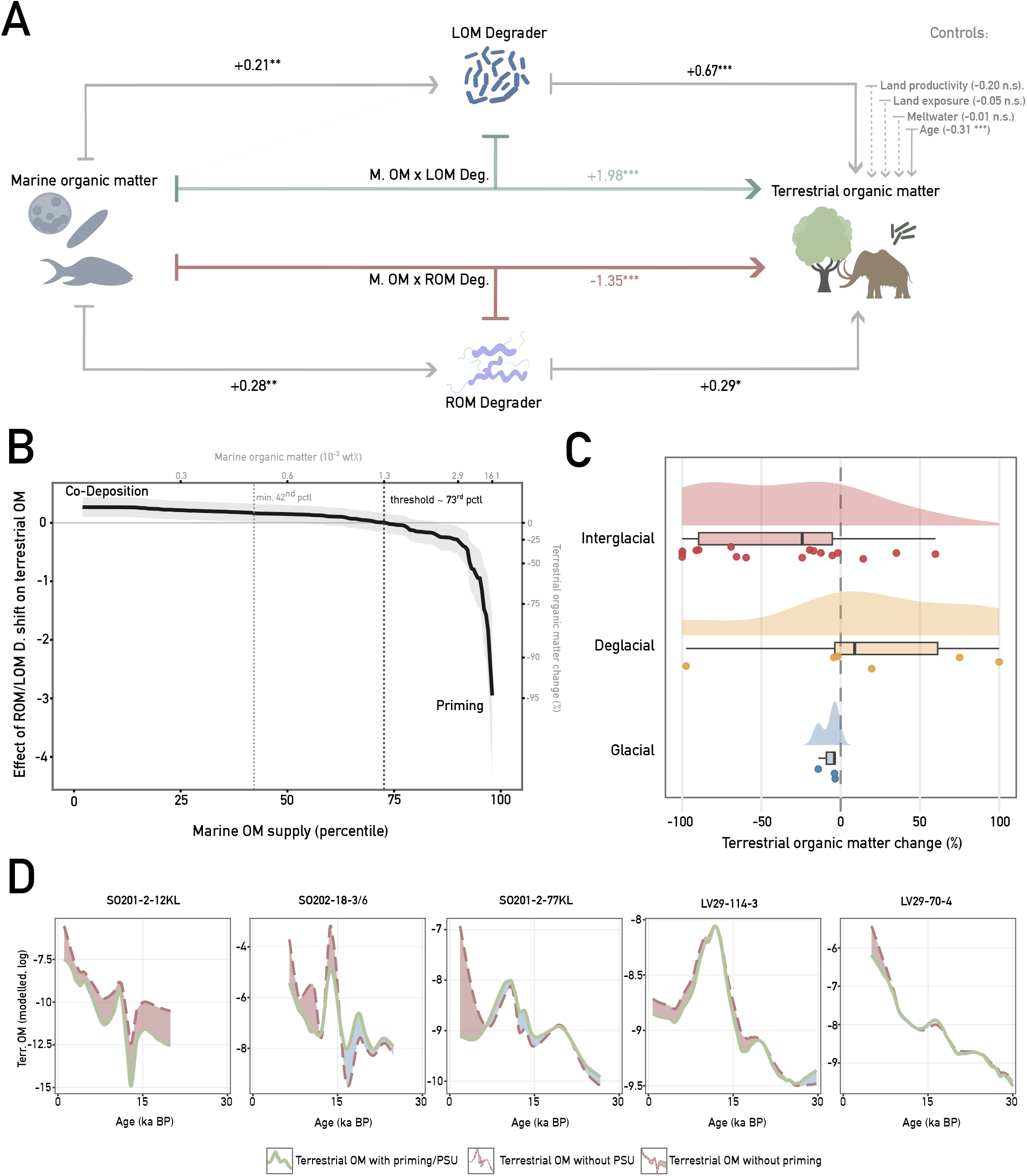
Priming and preferential substrate utilization (PSU) as processes controlled by marine organic matter and time. **A** Piecewise structural equation model (fitted as a Gamma(log) linear mixed model with core as a random effect) linking marine organic matter, labile-(LOM) and refractory-organic-matter (ROM) degrader biomass, and buried terrestrial organic matter. Arrows are standardized path coefficients: direct paths (marOM → LOM-deg. +0.21; marOM → ROM-deg. +0.28; LOM-deg. → terrOM +0.67; ROM-deg. → terrOM +0.29) and the two interaction paths — marOM × LOM-deg. (+1.98; preferential substrate utilization/preservation, light green) and marOM × ROM-deg. (−1.35; priming, light red). Grey dashed arrows are covariates (land productivity −0.20, land exposure −0.05, meltwater −0.01, age −0.31). *P<0.05, **P<0.01, ***P<0.001; n.s; marginal R^2^ = 0.73). **B–D, all derived from the model in A. B** Conditional (Johnson–Neyman) effect of a shift toward ROM-degrader dominance on terrestrial organic matter as a function of marine-organic-matter supply (95% band). Supply is shown as a percentile (lower axis) and as the corresponding absolute content (upper axis, 10□^3^ wt %), and the effect is expressed in model units (change in log terrestrial organic matter; left axis) as well as the percentage change in terrestrial organic matter per one-standard-deviation shift toward ROM-degrader dominance (right axis). The effect is near-zero to positive at low marOM (co-deposition) and turns significantly negative (priming) above the ~73rd percentile (≈1.3 × 10□^3^ wt %; dotted lines mark the 42nd and 73rd percentiles). **C** Model-attributed net change in terrestrial organic matter (%) by climate state (raincloud: density, box, samples). **D** Down-core terrestrial organic matter (modelled, log) for the five cores: full model (with priming and PSU; green) versus counterfactuals with priming or PSU removed (dashed); shaded areas quantify terrestrial OM preserved by PSU and lost to priming.

With physical delivery proxies showing no significant effect, terrestrial carbon dynamics appear primarily biologically driven (**Supplementary Figure 3**). Rather than acting through a single mechanism, LOM and ROM degraders independently shift the terrestrial pool away from the shared co-deposition baseline in opposite directions (**Fig. 3**). This baseline represents the normal state dictated by bulk sedimentation rates, where terrestrial and marine fractions of the bulk organic matter passively co-accumulate independent to any selective microbial alteration. As indicated by their positive interaction term, the simultaneous presence of marine organic matter and LOM-degraders results in a higher proportion of terrestrial organic matter than bulk co-deposition alone would predict. This pattern seems to reflect preferential substrate utilization. Much like in soil studies where microbes favour labile sugars over heavier compounds like lignin (Blagodatskaya et al., 2011), the marine community here appears to prioritize labile marine inputs. Such a metabolic shift would mechanistically explain how marine organic matter shields terrestrial carbon as originally proposed by Hedges et al. (1997). Conversely, the negative interaction between marine organic matter and ROM-degraders drives terrestrial organic matter below the co-deposition baseline. Therefore, our results provide evidence for a marine priming effect, demonstrating that labile marine inputs actively fuel the degradation of the more refractory terrestrial pool, providing a crucial missing link between short-term microbial interactions and millennial-scale carbon burial. We reveal that the ultimate fate of this terrestrial carbon is not uniformly dictated by marine organic matter availability, but is fundamentally governed by which microbial degrader group dominates (**Fig. 3A**). Our findings lead us to conceptualize priming and preferential substrate utilization not as opposing ends of a single spectrum, but as independent deviations from a shared co-deposition baseline. From this perspective preferential substrate utilization operates as a distinct LOM degrader mechanism rather than the direct inverse of priming. Viewing them as independent strategies implies that both can take place at the same time.

Our results show that priming and preferential substrate utilization are strictly regulated by organic matter availability (**Fig. 3B**). At low marine organic matter levels, the specific degrader group has only a small influence. This condition defines the co-deposition baseline, where terrestrial and marine carbon are simply co-deposited and preserved together (**Fig. 3B**). The microbial interaction only becomes significant once the marine substrate exceeds the ~73rd percentile of marine OC_DNA-projected_ (wt%) (~13 μg). This marks the threshold where the partial effect crosses from the co-deposition baseline to priming. Abundant labile marine organic matter is therefore a necessary condition for high remineralization rates.

### Temporal dynamics of priming and preferential substrate utilization

The microbial processing of terrestrial organic matter shifts from co-deposition during the glacial to active, priming-driven remineralization of terrestrial organic matter during the Holocene (Kruskal-Wallis over all intervals p = 0.009, ordered temporal trend p = 0.021, **Fig. 3C**). During glacial and deglacial intervals, marine organic matter rarely exceeds the necessary threshold (~73rd percentile of marine OC_DNA-projected_ wt%). However, when this threshold is surpassed, a higher biomass of LOM degraders establishes, driving preferential substrate utilization and consequently promoting the retention of terrestrial organic matter, most evident during the deglacial (**Fig. 3C**). During the interglacial, however, marine organic matter crosses the threshold more often and coincides with a ROM-degrader dominated bacterial community. This coupling causes net loss and removes a median of ~24 % of terrestrial organic matter where it is active (~13 % integrated across all interglacial samples). The system thus transitions over time, with the dominant process shifting from terrestrial carbon burial through co-deposition in the glacial, to preferential substrate utilization during the deglacial, and ultimately to priming-driven degradation within the interglacial. Projected onto the individual down-core records, these opposing effects resolve into discrete intervals of terrestrial carbon net negative (priming) and net positive (preferential substrate utilization) change relative to the co-deposition baseline (**Fig. 3D**).

For each core, we compare the modelled terrestrial organic matter under the full interaction (green) with counterfactuals where priming (red) or preferential substrate utilization (blue) is removed. Where the “without priming” counterfactual rises above the modelled trajectory (red line), priming actively remineralizes terrestrial carbon (**Fig. 3D**). This loss is concentrated in the substrate-rich, interglacial sections of cores characterized by a generally higher marine organic matter content (SO201-2-77KL, LV29-114-3, and SO202-18-3/6). By contrast, the “without PSU” (without preferential substrate utilization) counterfactual lies below the modelled trajectory (blue envelope) mainly in the older, low-substrate intervals (**Fig. 3D**). In these sections, preferential substrate utilization added to terrestrial preservation. In the marine organic matter-poor record LV29-70-4, the three trajectories nearly coincide, indicating only minimal microbial processing. The timing and magnitude of the two envelopes differ slightly among cores according to substrate availability. This pattern mirrors both the substrate threshold (**Fig. 3B**) and how the interactions differ between cold glacial and warm interglacial states (**Fig. 3C**).

The temporal trends indicate that both processes, priming and preferential substrate utilization are sensitive to environmental factors, in addition to microbial community structure and marine organic matter supply. To explicitly test for a temperature dependency, we incorporated the NGRIP □^1^□O temperature record (Rasmussen et al., 2006) into a separate structure equation model (**Supplementary Figure 4**). Temperature acted indirectly, structuring the algal community and through it the microbial degrader community composition and both processes. Warming increases marine organic matter availability (□ = +0.40) and strengthens the priming interaction (interaction □=−0.12*, per +1 SD temperature, equals roughly ~2–3‰ □^1^□O, **Supplementary Figure 4**) in all cores. Beyond this indirect pathway, temperature and other abiotic factors exert a first-order control on the terrestrial organic matter delivered to the marine realm. The composition and condition of the delivered terrestrial organic matter are especially sensitive to terrestrial abiotic changes and fluvial transport dynamics and are therefore impacted by shifting climate (Winterfeld et al., 2018; Feng et al., 2013; Cao et al., 2023; Herzschuh et al., 2025, 2026). On land, glacial–interglacial shifts lead to changes in biomass production, river discharge, and coastal erosion, thereby determining the amount and reactivity of the initial terrestrial carbon input (Herzschuh et al., 2025, 2026).

**Figure 4.**
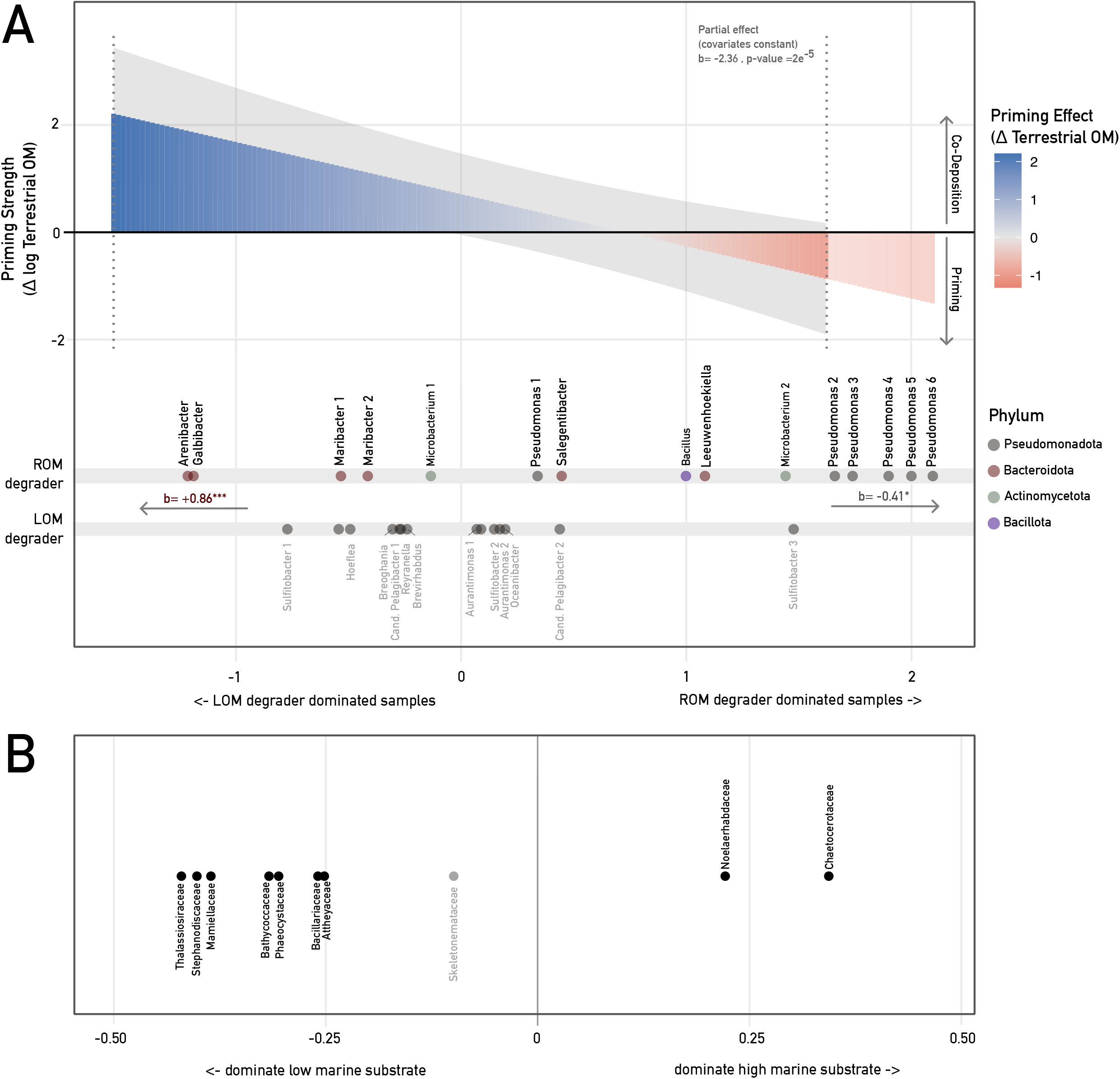
Taxonomic composition of the marOM-dependent shift between terrestrial organic matter priming and preferential substrate utilization. **A** Partial effect of the interaction between marine organic matter and degrader-community balance on buried terrestrial organic matter. Top: Modelled effect of interaction term on terrestrial OM (priming strength, □ log terrestrial OM; colour scale) along the degrader-balance gradient (x-axis), with covariates (total degrader organic matter, age, core) held constant. The balance axis represents the log-ratio of refractory-OM (ROM) to labile-OM (LOM) degrader biomass (left: LOM-degrader dominated; right: ROM-degrader dominated). Positive values (blue) indicate net terrestrial-OM gain via co-deposition; negative values (red) indicate net loss (priming). The grey envelope shows the 95% confidence band, and dotted lines delimit the observed data range. The substrate × degrader interaction is significant (b = −2.36, P = 2 × 10□□). Bottom: Bacterial degrader taxa ordered by their abundance-weighted position along the balance axis, grouped by degrader group and coloured by phylum. Within the ROM degrader group, the substrate interaction is phylum-specific: positive for Bacteroidota (b = +0.86, P < 0.001; preservation-like) and negative for Pseudomonadota (b = −0.41, P < 0.05; priming). Here, b denotes the partial regression coefficient. **B** Association of phytoplankton (marine primary-producer) families with marine-organic matter content over all cores, quantified as the Spearman rank correlation (□) between each family’s relative abundance and marine organic-matter content (x axis). Significant correlations are indicated by black dots, whereas non-significant ones are shown in grey. Families to the right dominate high-substrate samples, whereas families to the left dominate low-substrate samples.

### Taxonomic controls on priming versus preferential substrate utilization

Our results demonstrate that the taxonomic composition of the microbial community governs whether the net outcome of the interaction results in terrestrial organic matter preservation or loss. Evaluating the partial effect of the ROM-to LOM-degrader ratio reveals a shift from a positive to a negative impact on terrestrial organic matter (b = −2.36, p = 2 × 10□□; **Fig. 4A**). This zero-crossing marks the exact switch between two opposing pathways: a positive effect indicating preferential substrate utilization, where LOM degraders bypass terrestrial carbon, and a negative effect representing active priming by ROM degraders. By ordering the samples along this ratio and placing each taxon at its abundance-weighted average position, we can resolve exactly which taxa anchor these distinct biogeochemical states (**Fig. 4B**).

Mapping the ROM-degrader genera along this axis shows that they cluster clearly by phylum, driven by whether they target marine or terrestrial carbon sources (**Fig. 4B**). Bacteroidota, like *Arenibacter, Galbibacter, Maribacter*, and *Salegentibacter* occupy the LOM-dominated, preserving end (b = +0.86, p < 0.001). Conversely, Pseudomonadota, dominated by *Pseudomonas* spp., occupy the ROM-dominated, priming end (b = −0.41, p < 0.05). Thus, a single functional group of ROM-degraders harbors two distinct phyla with reversed impacts on terrestrial carbon storage. As demonstrated in coastal systems, members of Bacteroidota degrade high-molecular-weight marine polymers like proteins, cellulose, and chitin through specialized outer-membrane transporters (Fernández-Gómez et al., 2013). They largely lack the machinery for refractory terrestrial compounds such as lignin and thus bypass and spare terrestrial organic matter (Fernández-Gómez et al., 2013; Peng et al., 2024). In contrast, *Pseudomonas* exploit labile phytoplankton exudates (Granum et al., 2002) and can co-metabolize for example lignin, driving terrestrial degradation (Peng et al., 2024).

The LOM-degrader group is taxonomically anchored within the Pseudomonadota and spreads broadly across the axis (**Fig. 4B**), where methylotrophs and labile OM specialists (such as *Methyloceanibacter, Candidatus* Pelagibacter, and most *Sulfitobacter*) occupy the labile, carbon-retaining end. This distribution is consistent with preferential substrate utilization. *Methyloceanibacter*, for example, is restricted to simple, low-molecular weighted substrates (C1–C3) (Vekeman et al., 2016). This genus leaves more complex terrestrial matter untouched and therefore does not impact its burial. A minority of taxa pivot toward the priming side. *Aurantimonas* has metabolic plasticity that lets it switch to complex substrates under non-limiting conditions (Castillo-Ilabaca et al., 2026). Together with *Hoeflea, Breoghania*, and *Sulfitobacter*, it dominates a single core (SO202-18-3/6) where the LOM-degrader interaction is itself negative.

Finally, the principal phytoplankton sources are identified by their positive association with samples rich in marine organic matter (**Fig. 4C**). These enriched samples are dominated by bloom-forming Chaetocerotaceae and Noelaerhabdaceae, whereas low-organic matter samples are characterized by Thalassiosiraceae, Skeletonemataceae, and Phaeocystaceae. Notably, the prevalence of Chaetocerotaceae- and Noelaerhabdaceae-dominated samples increases significantly during the Holocene, a trend that our piecewise structural equation model identifies as being primarily driven by temperature. The temporal pattern of the phytoplankton community composition is consistent with previously reported ecological reconstructions from subarctic North Pacific sedaDNA records (Zimmermann et al., 2023; Buchwald et al., 2024; Dinkel et al., 2026).

Overall, these patterns demonstrate that microbial community composition, in the presence of marine organic matter, shapes the direction of carbon interactions. Depending on which metabolic strategies dominate, the marine organic matter fraction either stimulates the breakdown of terrestrial organic matter through priming or promotes its burial via preferential substrate utilization. In this way, shifts in microbial community structure directly regulate the efficiency of long-term terrestrial carbon storage on millennial timescale.

## Conclusion

We show that the fate of terrestrial carbon is dictated not only by its intrinsic chemical stability, but by a dynamic microbial filter sensitive to the amount of marine organic matter. Shifting the focus from chemical recalcitrance to ecological interactions reveals that the interplay between marine organic matter and specialized microbial degraders acts as a decisive switch. This switch either accelerates terrestrial carbon remineralisation through priming or facilitates its long-term burial via preferential substrate utilization (PSU). Crucially, the labile marine organic matter required for either priming or preferential substrate utilization is supplied by warming-induced phytoplankton blooms. Warming increases marine organic matter availability and strengthens especially the priming interaction in the North Pacific. The preservation of buried terrestrial carbon is therefore not a fixed molecular property but a climate-sensitive outcome of microbial ecology. This raises the possibility that continued warming shifts these processes toward net terrestrial carbon loss, turning a long-term sink into a source. Testing that feedback requires measurements beyond sedaDNA, but the temperature sensitivity we observe identifies the mechanism that may drive it.

## Material and methods

### Marine sediment cores

The sediment cores analysed for this study are all located in the Northern Pacific. The material of the northernmost site SO202-18-3/6 was recovered during R/V Sonne cruise SO202-INOPEX in 2009 (Gersonde, 2012) close to the continental shelf in the Bering Sea. This site consists of two cores, Piston Core SO202-18-3 (60.13° N, 179.44° W, water depth: 1111 m) and neighboring Kasten Core SO202-18-6 (60.13° N, 179.44° W, water depth: 1107 m) that were spliced into a composite record (Kuehn et al., 2014). The other two cores from the Bering Sea are SO201-2-77KL, located on the Shirsov Ridge (56.33°N, 170.69°E), and core SO201-2-12KL (off Kamchatka, 53.99°N, 162.37°E), both were recovered during Sonne cruise SO201-KALMAR Leg 2 in 2009. The next sediment cores LV29-114-3 (49,37° N, 152,87° E) and LV29-70-4 (48,00° N, 146,14° E) are located in the Okhotsk Sea and were recovered during Cruise LV29 with R/V Akademik M.A. Lavrentiev in 2002 (Biebow et al., 2002).

### Laboratory analysis

The extraction of sedimentary ancient DNA and the preparation of the shotgun library were carried out in paleogenetic laboratories at the Alfred Wegener Institute (AWI) Helmholtz Centre for Polar and Marine Research in Potsdam, Germany. The laboratory process for cores SO201-2-12KL and SO201-2-77KL is described in Zimmermann et al., 2023 and Buchwald et al., 2024, respectively. For this study 3 new marine sediment cores from the Northern Pacific were used.

We extracted DNA from all sediment samples leveraging the DNeasy PowerMax Soil Kit (Qiagen) and refined the extracts using the GeneJET PCR Purification Kit (Thermo Fisher Scientific). The workflow was organized into batches of nine samples plus one negative control (EB) to validate the extraction chemicals and handling procedures. After quantifying the DNA with a Qubit Fluorometer (Thermo Fisher Scientific), we adjusted each extract to a standardized concentration of 3 ng μL□^1^ for downstream analysis.

Single-stranded DNA libraries were constructed following the protocol of Gansauge et al., 2017, incorporating modifications specified by Schulte et al., 2021. Following initial library construction, quantitative PCR (qPCR) was utilized to determine the optimal number of amplification cycles for subsequent index PCR. Sample libraries were amplified for 11–13 cycles, while control blanks were restricted to 11 cycles. This indexing step utilized P5 and P7 indexing primers (IDT) and AccuPrime Pfx polymerase (Life Technologies) with a starting library volume of 24 μl. Post-amplification, sample libraries were pooled at equimolar concentrations. To account for low-input controls, extraction and library blanks were included in the pool with a fixed volume of 1 μl each. The final library pool was purified and concentrated to a target volume of 10 μl and a molarity of 20 nM using the MinElute PCR Purification Kit (QIAGEN). Sequencing was performed on an Illumina NextSeq 2000 platform using P3 Reagents for 200 cycles, ensuring high-depth taxonomic coverage across the sedimentary record.

### Bioinformatic analysis

For bioinformatic analysis of the raw sequencing data, the alignment based HOLI pipeline was used (Pedersen et al., 2016), modified by Liu et al., 2024 with a larger customized taxonomic reference database and implemented in Nextflow (v23.10.0) for this study. The computational workflow starts with quality control and the removal of duplicate sequences to ensure data integrity. High-quality reads, pre-filtered via Fastp to a minimum effective length of 30 bp, were mapped against the RefSeq and custom reference databases using Bowtie2. To prioritize full-length sequence fidelity and avoid spurious local matches, the aligner was deployed in end-to-end mode. For the RefSeq alignment specifically, we implemented a retention threshold of up to 1,000 unique and valid alignments per read (-k 1000).

Taxonomic assignment was executed using the ngsLCA (v1.0.5) software, applying a stringent 95% minimum identity threshold. For custom databases, this classification was facilitated via metaDMG, which integrates the ngsLCA algorithm into its internal processing architecture. Following classification, taxonomically assigned reads were aggregated into comprehensive count matrices. These biological data were then synchronized with geochronological and stratigraphic metadata (sample depth and age) to produce a unified master dataset. All subsequent statistical modeling and downstream ecological analyses were conducted within the R environment (v4.4.2).

### Data Mining and Automated Habitat Assignment

Taxonomic classification of DNA sequence reads was performed to at least the Superkingdom level (Bacteria, Archaea and Eukaryota). To address the challenges of high-resolution habitat classification for all classified Bacteria and Archaea, we utilized MICROSPyDER (MICRObial Source crawling & Integration Python Data Engine for Relative-affinities), following the framework established by Yong & Weiß et al. (in press, 2026). This integrated pipeline utilizes a custom web crawler (Selenium and BeautifulSoup4) to systematically retrieve structured metadata from the MicrobeAtlas database (Rodrigues et al., 2017; Rodrigues et al., 2026). A Habitat Affinity Score (*H*_*i*_) was calculated to place each taxon *i* along a continuous spectrum. For Metazoa, Viridiplantae, and Fungi, we applied automated literature research using the regex package in R (v4.4.2), followed by a decision-tree determination of the origin habitat and manual verification.

### Organic carbon estimation - genC pipeline

To translate taxonomic read abundances into ecologically meaningful units, we employed the genC pipeline (genome-based Carbon reconstruction), as previously described by Yong & Weiß et al. (in press, 2026). The pipeline bridges the gap between physical laboratory measurements and biological reality through a two-stage reconstruction model. The pipeline was run, using the same thresholds and structural factors, as well as the same databases. Finally the same stress test (perturbation analysis) and the same sensitivity analysis (Sobol) were applied.

### Clustering for statistical model

First, to achieve a comprehensive functional profiling of the microbial community, we extended the classification of Labile Organic Matter (LOM) and Recalcitrant Organic Matter (ROM) degraders beyond the detected taxa in our samples. We performed a systematic screening of all bacterial and archaeal species retrieved during the global data mining process (see *MICROSPyDER pipeline*). Using string-matching algorithms (Regular Expressions), we queried the scientific names of all species against established functional patterns (Louca et al., 2016). LOM degraders were identified using patterns targeting taxa known for the rapid turnover of fresh organic matter (e.g., Vibrio, Pseudoalteromonas, Flavobacterium). Conversely, ROM-degraders were identified via patterns for taxa specialized in the breakdown of complex, soil-derived, or woody compounds, including *Streptomyces, Sphingomonas*, and *Caulobacter*. This systematic screening allowed us to calculate the relative read proportions of each functional group within the total bacterial community across all samples (Supplementary Data 2).

The marine LOM pool was defined as the sum of organic matter from aquatic algae and aquatic Metazoa. The terrestrial organic matter pool was calculated as the sum of Viridiplantae and terrestrial Metazoa.

### Model Selection and Evaluation of interaction of carbon pools

We modelled the fate of terrestrial organic carbon as a function of the marine microbial degrader community and the availability of labile marine substrate. The response variable was terrestrial organic matter (calculated via genC). Because terrestrial organic matter is strictly positive and strongly right-skewed with variance increasing with the mean, we used a generalized linear mixed model (GLMM) with a Gamma family and log link (lme4::glmer). A Gaussian model on log-transformed organic matter provided equivalent inference and was used for the partial-effect visualisations.

Community composition was expressed as the guild balance. We defined this as the z-standardised log-ratio of refractory to labile organic matter degrader organic matter (log(ROM/LOM)). Total degrader load (z(log(ROM + LOM))) was retained as a covariate to de-collinearise the balance from overall abundance. Priming was specified as the interaction between labile marine substrate and guild balance (substrate × log(ROM/LOM)). In a second, taxon-explicit parameterisation, we replaced the balance with the two individual guild abundances and their separate interactions with substrate (substrate × ROM-degrader and substrate × LOM-degrader). This isolated priming and preferential substrate utilisation (PSU) as distinct terms. All continuous predictors were z-standardised. Sample age (z-standardised) was included as a covariate, and core identity was added as a random intercept to account for non-independence within the five cores.

Model selection relied on likelihood-ratio tests for nested models alongside the change in marginal and conditional R^2^ (MuMIn::r.squaredGLMM). We first tested whether terrestrial burial was controlled by terrestrial supply. To do this, we added a delivery block to a co-deposition baseline model (substrate + total load + age + core). This delivery block included meltwater discharge, deltaic land-area change, and terrestrial productivity, which were log-transformed due to strong right-skewness. The delivery block added < 2 % of explained variance and did not meaningfully improve the fit (LRT p ≈ 0.05). Its only non-zero term, land productivity, showed a sign reversal indicative of collinearity. By contrast, the substrate × balance interaction was retained in every specification (□ ≈ −2.0, p < 10□^3^; log-response partial-effect model b = −2.36, p = 2 × 10□□).

To evaluate the interaction, we mapped the marginal effect of the guild balance on terrestrial organic matter across the observed range of marine substrate using a Johnson-Neyman analysis. The effect is significantly positive (indicating co-deposition) below the ~46th substrate percentile. It crosses zero at the ~73rd percentile and becomes significantly negative (indicating priming) above the ~85th percentile. Partial-effect conditioning curves were obtained by predicting terrestrial organic matter across the balance gradient at fixed low and high substrate levels, with all covariates held at their means (Fig. 3).

For the taxonomic resolution (Fig. 4), each degrader genus was positioned along the same guild-balance axis at its abundance-weighted mean balance. This represents a weighted-average score rather than an ordination axis. Phytoplankton families were positioned based on their Spearman rank correlation with marine substrate. Within the ROM-degrader guild, we tested whether phyla differed in their effects. We regressed terrestrial organic matter on the substrate interactions of the z-standardised summed relative abundances of ROM-associated Pseudomonadota and Bacteroidota.

### Statistics

All analyses were performed in R (v. 4.5.1). Mixed models were fitted using lme4 (Gamma GLMM) and nlme (Gaussian LMM within the structural equation model). Marginal and conditional R^2^ values were obtained using MuMIn. Continuous variables were z-standardised prior to modelling so that coefficients of standardised predictors act as standardised effect sizes. Fixed-effect significance was assessed with Wald tests, and nested models were compared using likelihood-ratio tests. Significance was set at □ = 0.05 (*** p < 0.001, p < 0.01, * p < 0.05).

To quantify the effect of the interaction on terrestrial carbon for each sample, we calculated the percentage change relative to a counterfactual model without the interaction term. This was calculated as (1 − exp(contribution)) × 100, where positive values denote loss (priming) and negative values denote gain (PSU), capped at ±100 %. Samples were assigned to specific climate epochs (glacial, deglacial, interglacial). The per-epoch distributions were non-normal and zero-inflated because most intervals were substrate-limited and therefore inactive.

Consequently, differences across epochs were tested over all intervals using the Kruskal-Wallis test. We also applied an ordered trend test (glacial to interglacial) based on the Spearman correlation between the per-sample effect and epoch rank, evaluated by 20,000 permutations. The frequency of priming-active intervals across epochs was assessed using Fisher’s exact test. Epoch medians are reported alongside interquartile ranges. Means over all intervals, with inactive samples set to zero, are reported as record-integrated losses.

The environmental control on the interaction was tested using a piecewise structural equation model (piecewiseSEM) with core as a random intercept (nlme::lme). Northern Hemisphere temperature was represented by NGRIP □^1^□O (Rasmussen et al., 2006), matched to each sample by the nearest age. The model specified marine substrate and community balance as functions of temperature. Terrestrial organic matter was modeled as a function of the substrate × balance interaction and age. Global fit was assessed using tests of directed separation and Fisher’s C, while standardised path coefficients were obtained by scaling. Temperature moderation of priming was evaluated via the three-way substrate × balance × temperature interaction in the terrestrial organic matter model.

