## Supplementary Material for "Microbial priming controls terrestrial carbon burial in marine sediments"

**Supplementary Information**

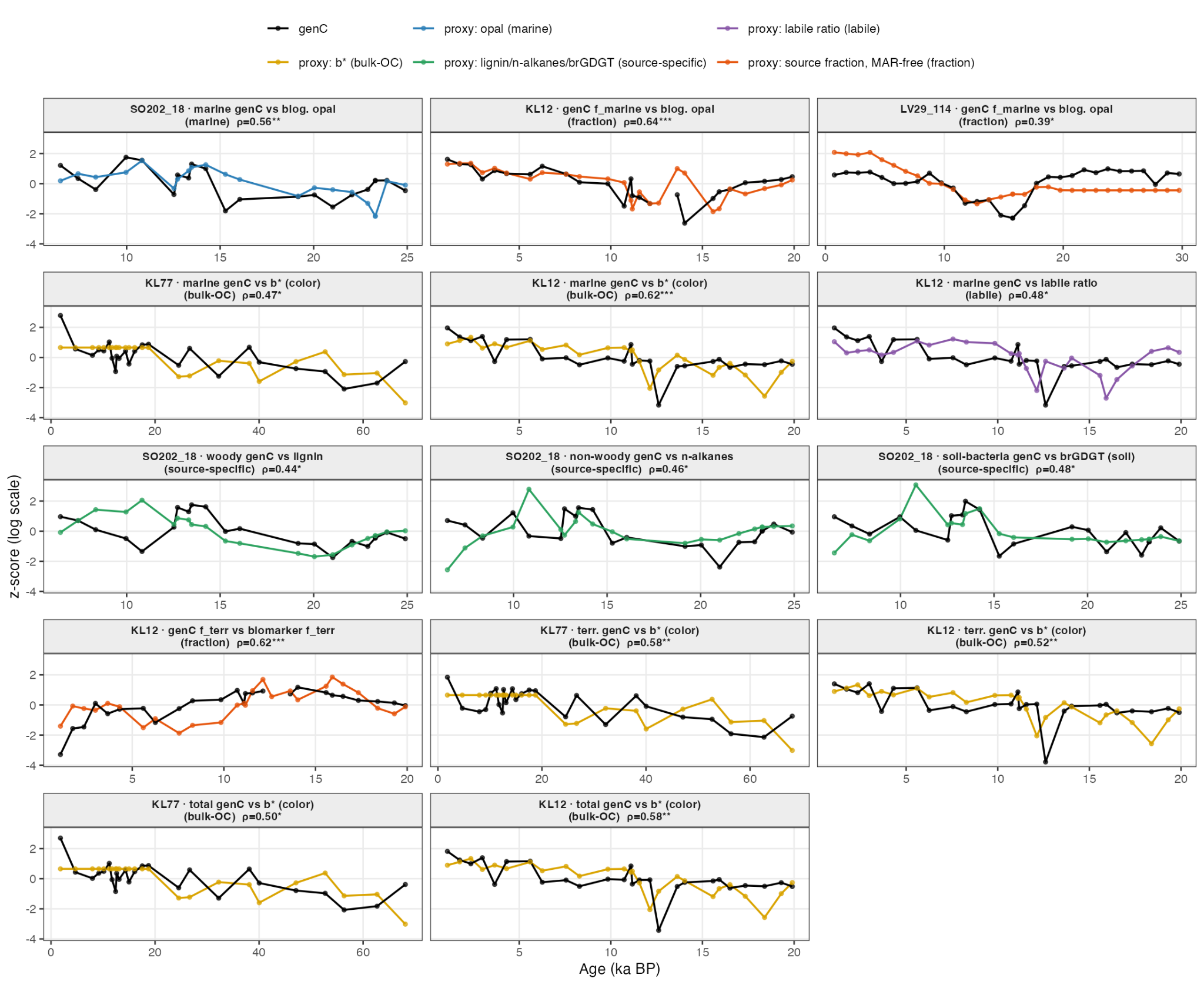

**Supplementary Figure 1. Validation of the DNA-based organic-carbon reconstructions (genC) against independent proxies.** Down-core comparison of genC (black) with established, independent organic-matter proxies (coloured) for the cores and organic-matter fractions where such proxies are available. In each panel both series are log-transformed and z-scored (no dual axis), and their agreement is quantified by the raw Spearman rank correlation (ρ; *P < 0.05, **P < 0.01, ***P < 0.001), computed without age-detrending because a valid proxy is expected to reproduce the down-core trend. Proxy types (legend): biogenic opal (marine productivity), sediment colour b* (bulk organic carbon), lignin phenols, long-chain n-alkanes and branched GDGTs (source-specific terrestrial), the labile fatty-acid ratio, and marine/terrestrial source fractions computed to be independent of the mass-accumulation rate (MAR-free). Panels are grouped by the genC component tested — marine genC (top rows), terrestrial genC (middle rows) and total genC (bottom row). Across all comparisons genC covaries positively and significantly with the independent proxies (ρ = 0.39–0.64), spanning marine (opal), bulk-OC (b*), source-specific terrestrial (lignin, n-alkanes, brGDGT) and composition-based (fraction) records, demonstrating that the sedaDNA-derived carbon pools track independent sedimentary indicators of organic-matter source and amount.+

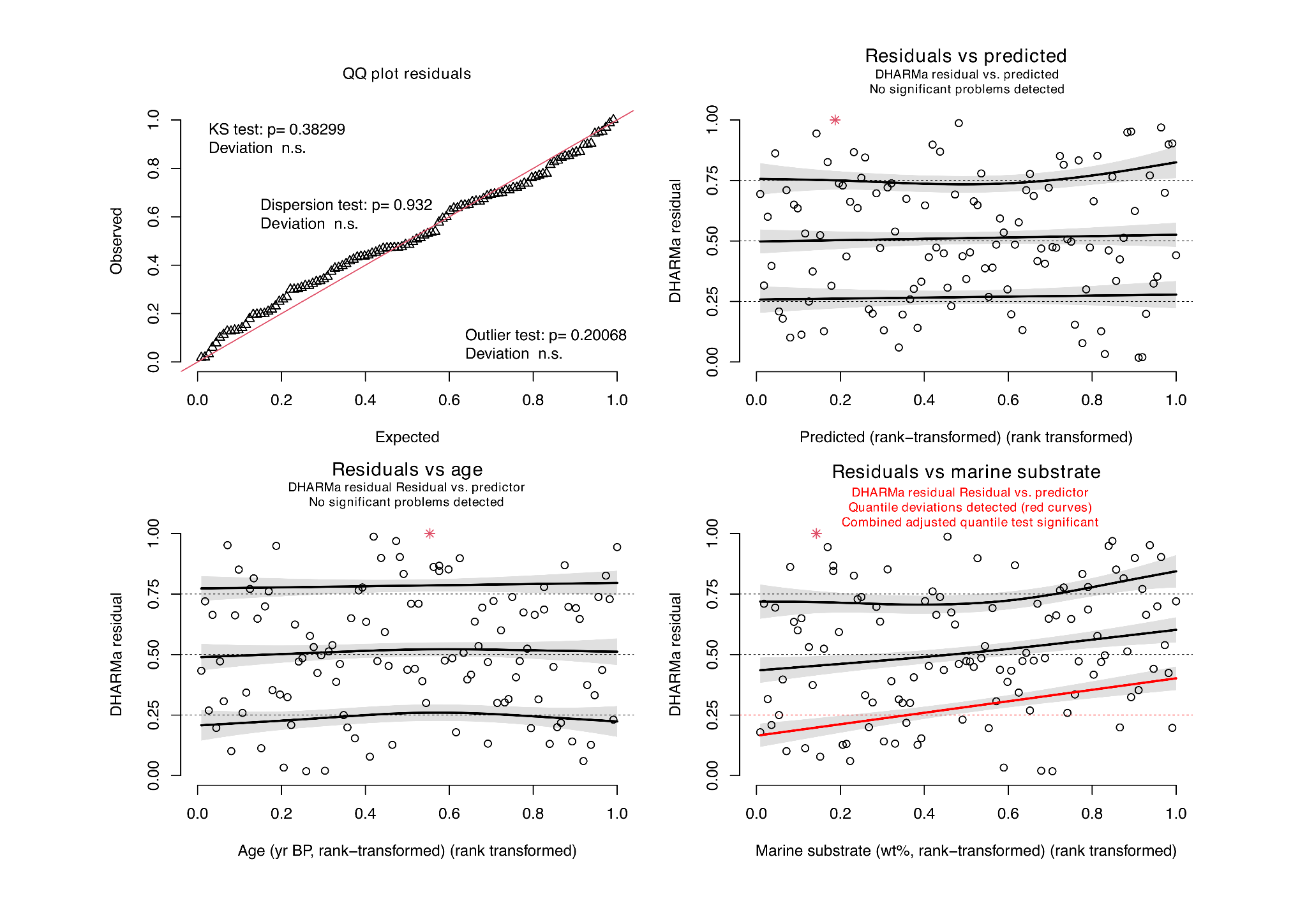

**Supplementary Figure 2. Residual diagnostics of the Gamma (log-link) mixed model.** Simulated, scaled quantile residuals of the fitted Gamma(log) GLMM (terrestrial organic matter ~ marine organic matter × ROM- and LOM-degraders + covariates + (1 | core)), computed with DHARMa (1,000 simulations). For a correctly specified model these residuals are uniformly distributed on [0, 1] and show no structure against predictors. Top left, quantile–quantile plot of the scaled residuals against the expected uniform distribution; the Kolmogorov–Smirnov uniformity test (P = 0.38), the dispersion test (P = 0.93) and the outlier test (P = 0.20) are all non-significant, indicating that the distributional, dispersion and outlier assumptions are met. Top right and bottom left, scaled residuals against the model predictions and against sample age; the 0.25, 0.5 and 0.75 quantile-regression lines (solid) are flat, showing no residual structure with fitted values or along the temporal axis (no unaccounted-for age trend). Bottom right, scaled residuals against marine substrate (marine organic matter); here the quantile lines deviate significantly (red), reflecting the non-linear, threshold-dependent effect of marine organic matter on terrestrial-carbon fate that the linear interaction term does not fully capture — the pattern that motivates the Johnson–Neyman threshold analysis (Fig. X).

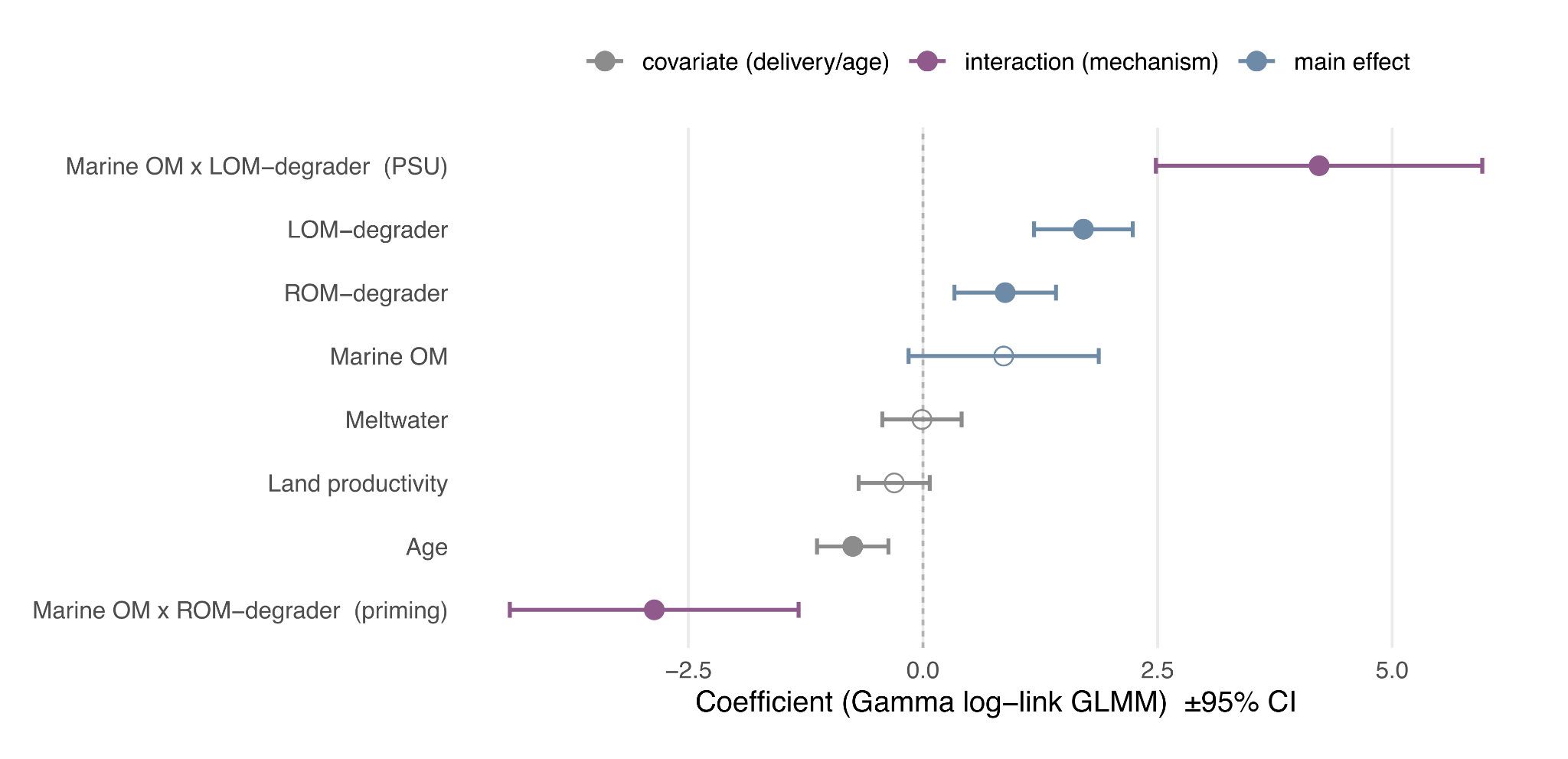

**Supplementary Figure 3. Coefficients of the Gamma (log-link) mixed model of terrestrial-carbon storage.** Estimated coefficients (± 95% confidence intervals) of the Gamma(log) generalized linear mixed model (terrestrial organic matter ~ marine OM × ROM-degraders + marine OM × LOM-degraders + covariates + (1 | core); n = 112). Predictors were z-scored and coefficients are on the log link, so effect sizes are comparable within the model. Symbols are coloured by term type — the two marine organic matter × degrader interactions (mechanism, purple), the degrader and marine organic matter main effects (blue), and the delivery and age covariates (grey) — and filled symbols denote significance (P < 0.05), open symbols non-significance. The two interaction terms dominate the model and have opposing signs: marine OM × LOM-degrader is strongly positive (preferential substrate utilization) and marine OM × ROM-degrader strongly negative (priming). In contrast, the terrestrial-delivery covariates (meltwater, land productivity) are non-significant.

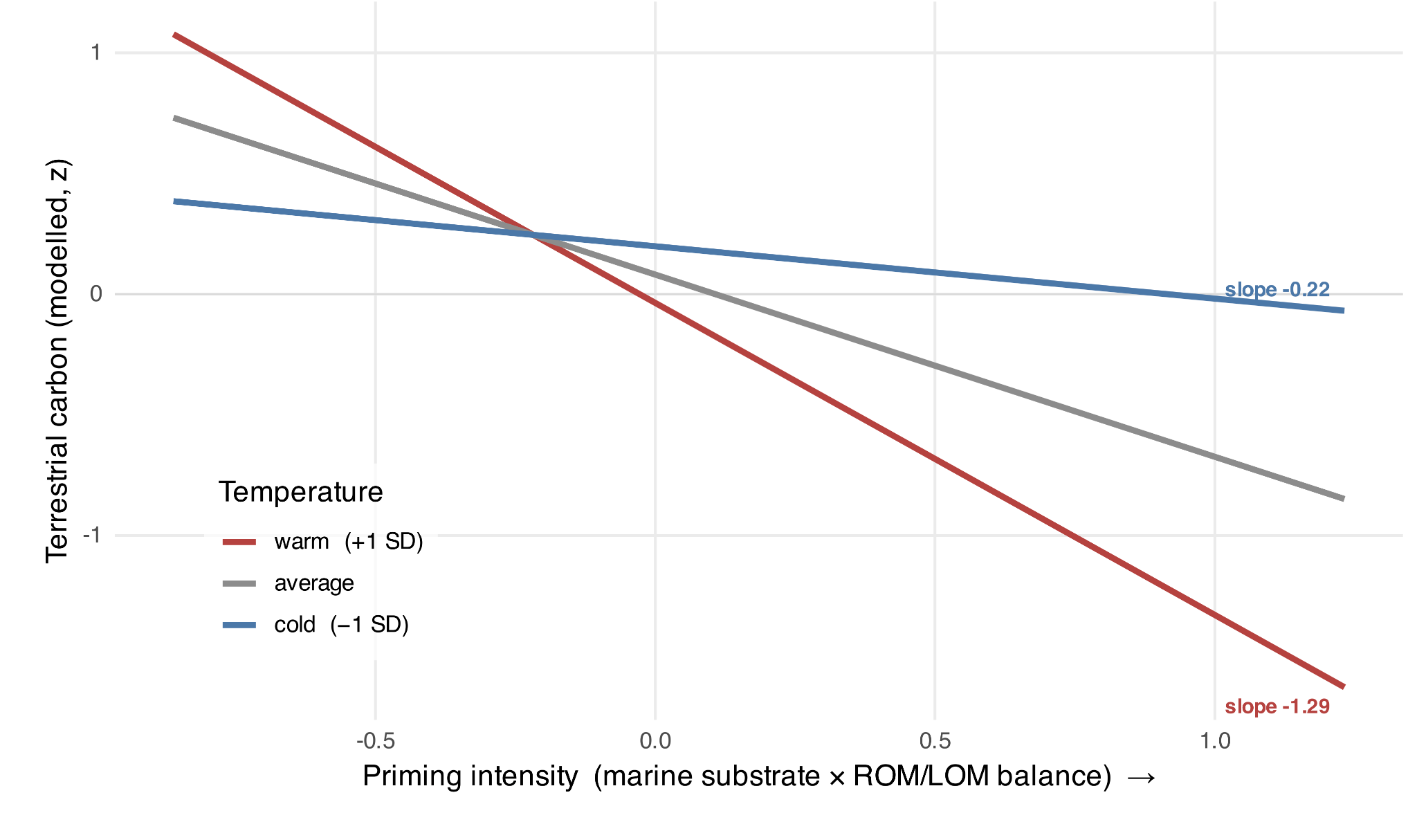

**Supplementary Figure 4. Warming steepens the priming effect on terrestrial carbon.** Simple slopes of the priming → terrestrial-carbon relationship, evaluated at cold (−1 SD, blue), average (grey) and warm (+1 SD, red) NGRIP temperature. The x-axis is priming intensity (marine substrate × ROM/LOM community balance, z-scored); the y-axis is modelled terrestrial carbon (z-scored, log scale), holding marine substrate and total degrader biomass at their means. The negative slope steepens from −0.22 under cold to −1.29 under warm conditions, i.e. a given amount of priming removes progressively more terrestrial carbon as the climate warms (temperature × priming interaction β = −0.12, P < 0.05). The lines fan out at high priming intensity and converge at low intensity, so temperature only matters where priming is already active. Predictions are from a linear model (terrestrial carbon ~ substrate + priming × temperature + total degrader biomass) fitted across all cores.

**Supplementary Table 1. Total shotgun reads per sediment core.**

| **Core ID** | **Location** | **Total Shotgun Reads** |
| --- | --- | --- |
| LV29-114-3 | Okhotsk Sea | 30,657,841 |
| SO202-18-3/6 | Bering Sea | 27,466,802 |
| LV29-70-4 | Okhotsk Sea | 25,272,676 |
| KL12 | Bering Sea | 13,133,754 |
| KL77 | Bering Sea | 8,549,227 |
